# Distinct HPO axis responses and ovarian aging trajectories to chronic unpredictable mild stress in reproductively young versus middle-aged female mice

**DOI:** 10.64898/2026.04.24.720585

**Authors:** Tiannuo Yang, Shuqin Zhang, Danqi Liu, Laijia Li, Kunpeng Zhou, Yufeng Han, Jianchuan Wang, Hanwen Zhang, Yiqing Ma, Shangxuan Liu, Boyu Ma, Furui Jin, Jian Li, Yudong Wang, Zelan Hu

**Author notes:** Correspondence: Jian Li, Yudong Wang, Zelan Hu. These authors contributed equally to this work (Tiannuo Yang, Shuqin Zhang, Danqi Liu).

## Abstract

Psychosocial stressors are key contributors to ovarian functional decline. Chronic unpredictable mild stress (CUMS) is widely used to model stress-induced premature ovarian insufficiency (POI) in mice; however, current animal models do not adequately reflect middle-aged women, who represent a key population exposed to chronic psychosocial stress, nor do they capture the dynamic progression toward POI. Here, female C57BL/6 mice aged 2 or 6 months were subjected to CUMS for 8 or 12 weeks. Estrous cyclicity, endocrine profiles, ovarian histology, and transcriptomic changes in HPO axis–related tissues were systematically analyzed. After 8 weeks of exposure, 2-month-old mice exhibited impaired pituitary responsiveness to estradiol negative feedback, as evidenced by dysregulated FSH secretion, indicating reduced stress tolerance compared with 6-month-old mice. Following 12 weeks of CUMS exposure, both age groups showed significant reductions in ovarian size and follicle numbers across all developmental stages. These findings demonstrate that CUMS induces an age-dependent progression toward POI, with short-term exposure eliciting compensatory phases preceding overt ovarian insufficiency, accompanied by distinct endocrine and reproductive alterations and differential responsiveness of the HPO axis. Transcriptomic analyses revealed age-dependent stress responses: ovaries of 2-month-old mice displayed marked activation of inflammatory and immune-related pathways, whereas 6-month-old mice showed sustained upregulation of protein kinase-related signaling networks. Notably, the 6-month-old CUMS model more closely recapitulates stress-associated reproductive aging in adult women.

**In brief:** CUMS has been widely used to establish mouse models of psychosocial stress–induced POI. However, current animal models do not adequately reflect middle-aged women, who represent a key population exposed to chronic psychosocial stress, nor do they capture the dynamic progression toward premature ovarian insufficiency (POI). In this study, we demonstrate that different durations of CUMS exposure induce distinct stages of ovarian dysfunction in both young and middle-aged mice, with short-term exposure driving age-dependent compensatory phases and prolonged exposure leading to overt POI, both accompanied by divergent endocrine and reproductive alterations, alongside age-dependent changes in HPO axis responsiveness to CUMS. Notably, the 6-month-old CUMS model shows greater clinical relevance in recapitulating chronic psychosocial stress and stress-related reproductive aging in adult women.

## Introduction

The ovary is the pivotal organ in female reproductive aging(1), and its functional decline is primarily characterized by a reduction in follicle quantity and a decline in oocyte quality(2). This decline is influenced by a combination of genetic, environmental, psychosocial, and lifestyle factors (1) and is associated with various comorbidities such as osteoporosis, cardiovascular diseases, and cognitive decline (3). Within the spectrum of ovarian functional decline, physiological aging and pathological conditions represent distinct trajectories. Accelerated or excessive follicle depletion can drive ovarian aging beyond its normal physiological limits, culminating in a pathological state known as premature ovarian insufficiency (POI).

Clinically, POI is defined as a pathological decline of ovarian function before the age of 40 and is characterized by menstrual disturbances, elevated follicle-stimulating hormone (FSH > 25 U/L), and fluctuating reductions in circulating estrogen levels (4). Increasing evidence highlights chronic psychosocial stress as a key environmental contributor to the pathophysiology of POI, primarily by perturbing the hypothalamic-pituitary-ovarian (HPO) axis. Specifically, sustained stress leads to persistent activation of the hypothalamic-pituitary-adrenal (HPA) axis, resulting in excessive glucocorticoid secretion. This, in turn, inhibits gonadotropin-releasing pathways and ovarian steroidogenesis, thereby suppressing overall HPO axis activity (5). Concurrently, prolonged exposure to stress elevates reactive oxygen species (ROS) generation within ovarian tissue and triggers pro-inflammatory signaling cascades the ovary (6,7). Collectively, these stress-induced alterations impair folliculogenesis and disrupt endocrine function, as reflected by aberrant levels of anti-Müllerian hormone (AMH) and estradiol (E2) (8).

To experimentally investigate the mechanistic link between chronic stress and ovarian aging, the chronic unpredictable mild stress (CUMS) model has been widely employed. CUMS mimics human psychosocial stress by applying prolonged, low-intensity and unpredictable stressors(9). In C57BL/6 mice, CUMS markedly reduces the numbers of primordial and developing follicles in ovaries and decreases circulating AMH and E₂ levels(10). These alterations are accompanied by increased ovarian oxidative stress, accelerated granulosa cell senescence, and mitochondrial dysfunction(11,12). Collectively, these findings establish CUMS as a commonly employed paradigm to establish mouse models of POI, enabling further investigation into its pathophysiology and evaluation of potential therapeutic interventions(13).

However, a critical limitation of existing CUMS-based POI studies is their predominant focus on young animals. Most studies employing the CUMS model utilize young mice aged 6-8 weeks, which roughly correspond to human adolescence (14,15). Although mice at this age exhibit relatively stable physiological systems, their reproductive status remains in the young stage and thus fails to adequately model the unique physiological characteristics and stress sensitivity associated with the reproductive aging transition in middle-aged women(16).The middle-aged period represents a critical phase of reproductive functional decline in females and is also associated with a high incidence of POI(17). Based on these definitions, we categorized 2-month-old mice as reproductively young and 6-month-old mice as reproductively middle-aged(14), in order to capture age-dependent differences in the response of the HPO axis to stress intervention and in the progression of ovarian aging. Furthermore, accumulating evidence suggests that mice at pubertal and mature stages may exhibit significantly different physiological responses and capacities for injury repair following stress exposure (11). Consequently, investigating the effects of CUMS on ovarian aging in 6-month-old mice will not only deepen our understanding of the mechanisms underlying stress-induced reproductive aging but also provide important experimental evidence for psychological health management and fertility preservation in contemporary women.

Based on this rationale, the present study aimed to examine the impact of CUMS on ovarian aging in both reproductively young and middle-aged female mice. To delineate the dynamic role of stress during the progression of ovarian aging, female C57BL/6 mice at both ages were subjected to 8-week and 12-week CUMS. Following the intervention, we evaluated estrous cyclicity, serum hormone levels (including AMH, E2, FSH and LH), and follicle counts across developmental stages. In addition, to elucidate how unpredictable stress reshapes transcriptional patterns along the HPO axis, transcriptome sequencing and differential expression analysis was performed on pituitary, and ovarian tissues from each experimental group.

## Methods

### 1. Animals

All procedures were approved by the Institutional Animal Care and Use Committee of the International Peace Maternity and Child Health Hospital (Approval No. GKDW-A-2024-54). Female SPF-grade C57BL/6J mice aged 2 or 6 months were obtained from Zhejiang Vital River Laboratory Animal Technology Co., Ltd. Mice were housed five per cage under standard laboratory conditions (12-h light/12-h dark cycle, 25 C, 55-65% humidity) with ad libitum access to food and water. After one week of acclimatization, mice in the experimental groups were subjected to the CUMS protocol, whereas control mice were maintained under the same conditions as during acclimation.

### 2. Chronic Unpredictable Mild Stress (CUMS)

After acclimation, mice with regular estrous cycles were selected and randomly assigned to control or CUMS groups (15-20 mice per group). The CUMS paradigm consisted of various stressors including noise disturbance, overnight illumination, strobe light, foot shock, food or water deprivation, social crowding, dirty cage, wet bedding, empty bedding, tail suspension, and restraint stress. Detailed procedures are listed in Table 1. Body weight was measured weekly.

**Table 1.** Specific operations of various pressure sources.

| Stressor | Specific Procedure |
| --- | --- |
| Noise disturbance | Expose mice to noise levels of 60-75 dB for 1-3 h. |
| Nighttime illumination | Provide illumination during the night to disrupt mice's sleep. |
| Strobe light exposure | Apply low-intensity strobe lighting (100-300 flashes/min). |
| Foot shock | Place mice in a Skinner box and deliver foot shocks at an intensity of 0.3-0.7 mA, with a frequency of one shock every 3 s, lasting 3-10 min. |
| Food/water deprivation | Remove food or water from each cage for 12-24 h. |
| Social crowding | House 8-10 mice per cage for 12-36 h. |
| Wet bedding | Pour 50-200 mL of water into each cage and maintain wet bedding for 2-12 h. |
| Dirty bedding | Reduce the frequency of bedding changes or replace clean bedding with used bedding from control mice. |
| No bedding | Remove bedding entirely for 12-24 h. |
| Restraint stress | Restrain mice in a closed but ventilated 50 mL EP tube for 1-8 h. |
| Tail suspension | Suspend mice by the tail using a tail-suspension apparatus for 10-30 min. |

After sacrifice, ovaries were carefully dissected to remove surrounding fat tissue, dried briefly, and weighed using an analytical balance. Ovarian index was calculated as:

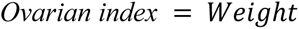

### 3. Estrous Cycle Assessment

Estrous cycles were monitored during the two weeks prior to the formal experiment and again during the final two weeks of the experimental period. The pre-experimental monitoring was performed to select female mice with regular estrous cycles for inclusion in the study, whereas the post-modeling monitoring was used to evaluate changes in estrous cyclicity induced by stress exposure. Vaginal cytology was performed daily at 8:00 a.m. for 14 consecutive days (covering approximately 2-3 estrous cycles). Vaginal lavage was conducted using a bent gavage needle to gently aspirate and expel approximately 0.2 mL of sterile physiological saline at the vaginal opening (inserted ∼2 cm). The collected fluid was placed onto a glass slide and air-dried at room temperature. Estrous stages were determined microscopically based on cell morphology: Proestrus: predominance of nucleated epithelial cells; Estrus: mainly anucleate cornified epithelial cells; Metestrus: mixture of cornified cells and neutrophils; Diestrus: predominance of leukocytes. Estrous cycle patterns were defined as: Normal cycle: 4-5 days with all four phases; Abnormal frequency: one phase abnormally prolonged or reduced; Cycle lengthened: >5 days. The latter two conditions were considered cycle irregularities.

### 4. Serum Hormone Measurements

Blood samples were centrifuged at 1500 r/min for 20 min, and serum was stored at -80 ℃ until use. Levels of E₂, FSH, AMH, LH, and CORT were quantified using commercial ELISA kits (Shanghai Ruifan Biotechnology Co., Ltd., Shanghai, China.). The assay sensitivities were as follows: E₂: 1.0 nmol/L; FSH: 10 pg/mL; AMH: 1.0 pmol/L; LH: 1.0 mIU/mL; CORT: 0.1 nmol/L.

### 5. Classification and Counting of Ovarian Follicles

Right ovaries (n = 5-6 per group) were fixed in 4% paraformaldehyde for at least 24 h, dehydrated, embedded in paraffin, and serially sectioned (3-5 μm). Every 20th section was selected for deparaffinization, rehydration, and HE staining. Follicles at different developmental stages were identified and counted according to established morphological criteria. Two independent investigators performed follicle quantification in a blinded manner.

### 6. RNA Sequencing of Mouse Tissues

For each tissue type, four cryovials were prepared, with each vial containing freshly dissected pituitary or ovarian tissue pooled from two mice. Samples were snap-frozen in liquid nitrogen and stored at -80 ℃. Total RNA was extracted from pituitary or ovarian followed by library preparation according to Illumina standard instruction (VAHTS Universal V6 RNA-seq Library Prep Kit for Illumina®). Agilent 4200 bioanalyzer was employed to evaluate the concentration and size distribution of cDNA library before sequencing with an Illumina novaseq6000. The protocol of high-throughput sequencing was fully according to the manufacturer’s instructions (Illumina). The raw reads were filtered by Seqtk before mapping to genome using Hisat2 (version:2.0.4). The fragments of genes were counted using stringtie (v1.3.3b) followed by TMM (trimmed mean of M values) normalization.

### 7. Statistical Analysis

Statistical analysis was conducted using GraphPad Prism software. All results are presented as mean ± standard deviation (SD). For two-group comparisons, unpaired Student’s t-tests were applied. For multiple group comparisons, one-way ANOVA was used. In vivo experiments were conducted using randomly assigned groups. No animals or data were excluded, and the study was not blinded; all data collection and analysis were objective.

### 8. Transcriptomic data analysis

Transcriptomic data analysis and visualization were performed in R (version ≥ 4.2.0). Gene expression matrices from ovarian and pituitary tissues were subjected to quality control and normalization, and lowly expressed genes were removed. Principal component analysis (PCA) was conducted using the normalized expression matrix, and samples were visualized based on the first two principal components. Differential expression analysis was performed using R-based statistical packages by comparing experimental groups or tissue types (ovary vs. pituitary). Genes with an absolute log_2_ fold change ≥ 0.25 and an adjusted P value < 0.05 were defined as differentially expressed genes (DEGs). Differential expression results were visualized using volcano plots, with log_2_ fold change on the x-axis and -log_10_(P value) on the y-axis. Gene Ontology (GO) enrichment analysis of DEGs was performed, including Biological Process, Cellular Component, and Molecular Function categories. GO terms with an adjusted P value < 0.05 were considered significant. Gene Set Enrichment Analysis (GSEA) was performed using ranked gene lists based on expression changes. Enrichment of the “gonadotropin-releasing hormone signaling pathway” and “response to estrogen” gene sets was evaluated using the normalized enrichment score (NES) and false discovery rate (FDR).

## Result

### 1. 8-week CUMS induced estrous cycle disruption in an age-independent manner

To better model the chronic social and life stress commonly experienced by women aged 30-50 years, we included a reproductively middle-aged group (6-month-old) alongside the conventional reproductively young group (2-month-old). Both groups were subjected to an 8-week CUMS paradigm (**Fig. 1A**) to systematically compare age-dependent physiological responses.

**Figure 1.**
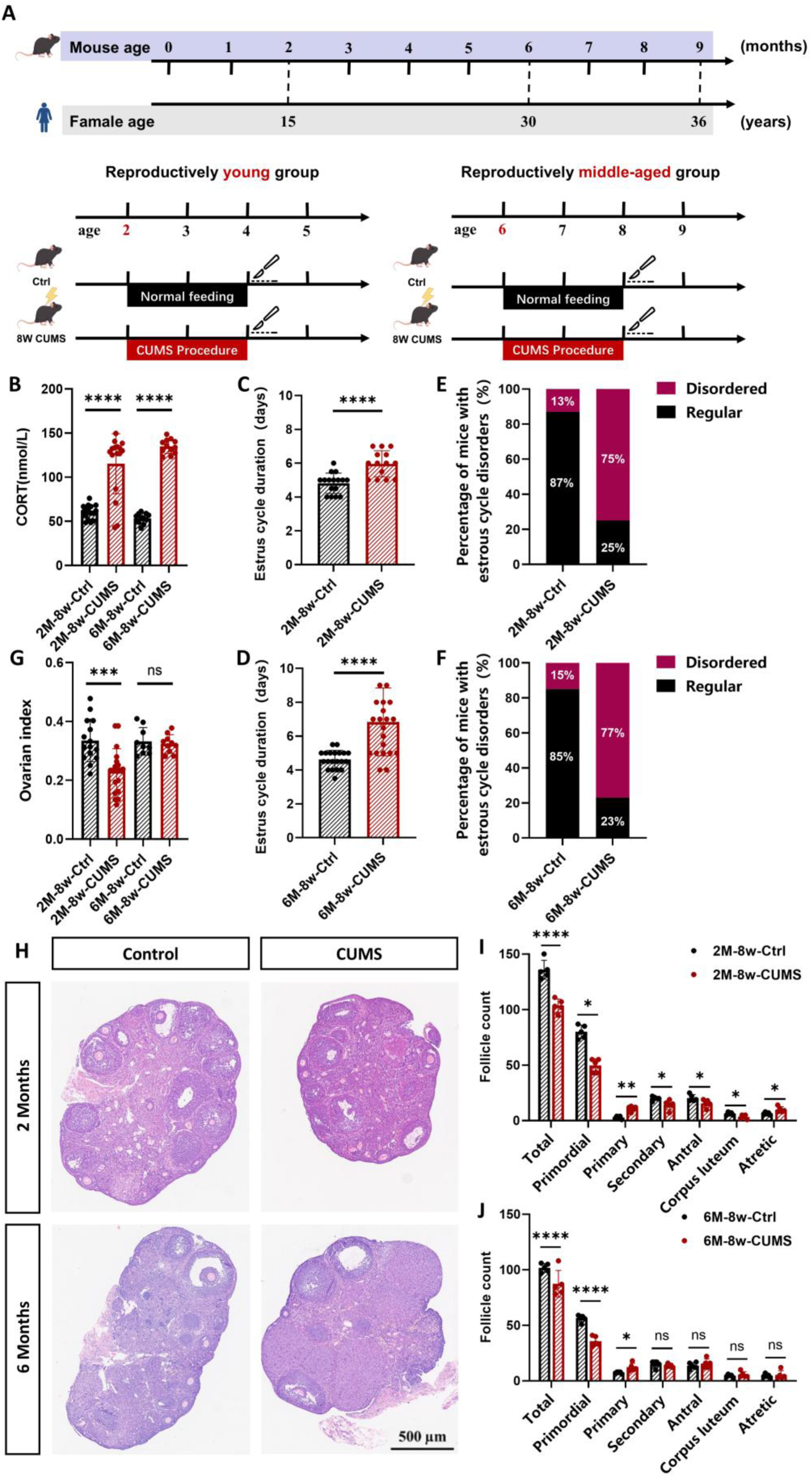
Effects of 8-week CUMS on estrous cyclicity and ovarian reserve in reproductively young and middle-aged mice. **(A)** Schematic diagram of the experimental design. **(B)** Serum corticosterone CORT levels in mice. **(C,D)** Estrus duration in the reproductively young group (C) and middle-aged group (D). **(E,F)** Proportion of mice with estrous cycle irregularities in the reproductively young group (E) and middle-aged group (F). **(G)** Ovarian index of mice. **(H)** HE staining of ovarian follicles. **(I,J)** Counts of different stages of follicles in the reproductively young group (I) and middle-aged group (J). Statistical analysis was performed using unpaired Student’s t-test for two-group comparisons, and one-way ANOVA followed by Tukey’s multiple comparisons test for multi-group comparisons. Data are presented as mean ± SEM. Statistical significance is indicated as ns not significant, * p < 0.05, ** p < 0.01, *** p < 0.001, **** p < 0.0001.

First, we validated the effectiveness of the stress model. Throughout the CUMS exposure, body weight remained consistently lower in both groups compared with their respective controls (**Fig. S1A, B**), indicating that chronic stress markedly inhibited normal weight gain. In parallel, serum corticosterone (CORT) levels were significantly elevated in both age groups (**Fig. 1B**), confirming successful activation of the sympathetic nervous system and the HPA axis.

To evaluate the impact of 8-week CUMS exposure on reproductive endocrine homeostasis, estrous cycles were examined daily for 14 consecutive days during the stress-induction period. In both age groups, CUMS treatment led to a pronounced lengthening of the overall estrous cycle compared with their respective controls **(Fig. 1C, D)**. Moreover, over 80% of exposed mice in both groups developed irregular cycles, as reflected by substantial increases in cycle-length variability **(Fig. 1E, F)**.

Collectively, the 8-week CUMS protocol effectively induced a physiological stress response and significantly disrupted estrous cyclicity in both young and middle-aged mice, characterized by prolonged and irregular cycles in the majority of animals.

### 2. 8-week CUMS induced ovarian atrophy and reserve depletion in an age-dependent manner

We then assessed the impact of chronic stress on ovarian health. In the reproductively young group, both ovarian weight (**Fig. S1C**) and ovarian index (**Fig. 1G**) were significantly reduced after the 8-week CUMS, indicating that this intervention alone was sufficient to induce marked ovarian atrophy. In contrast, no significant changes in these parameters were observed in the reproductively middle-aged group under the same stress paradigm (**Fig. S1C, 1G**).

Following the assessment of cycle disruption, we next examined whether CUMS affects ovarian follicular reserve. Histological examination with H&E staining revealed stress-induced alterations in folliculogenesis **(Fig. 1H)**. In both age groups, CUMS exposure resulted in a significant reduction in primordial follicles, accompanied by an increase in primary follicles, and an overall decrease in total follicle number **(Fig. 1I, J)**. These findings indicate that chronic stress accelerates the recruitment and depletion of the primordial follicle pool, leading to the buildup of early-stage follicles with impaired developmental progression. Notably, age-dependent differences were observed in advanced follicular stages. In the reproductively young group, the numbers of secondary follicles, antral follicles, and corpora lutea were significantly reduced, while the number of atretic follicles was significantly increased **(Fig. 1I)**. In contrast, no significant changes in these parameters were detected in the reproductively middle-aged group **(Fig. 1J)**. This differential pattern aligns with our earlier observation that 8-week CUMS induces ovarian atrophy in reproductively young mice, whereas middle-aged mice exhibit greater ovarian resilience.

Taken together, these results demonstrate that 8-week CUMS initiated premature follicle recruitment in both age groups. However, progression to advanced follicular depletion and ovarian atrophy occurred exclusively in reproductively young mice, underscoring a resilience to stress-induced ovarian failure in middle-aged animals.

### 3. Age-dependent HPO axis hormone responses to 8-week CUMS

To further investigate age-dependent responses of reproductively young and middle-aged mice to CUMS, we examined key hormones along the HPO axis. Both reproductively young and middle-aged group mice exhibited significant increases in AMH, E2 and LH levels **(Fig. 2A-C)**. These increase, together with accelerated transition from primordial to primary follicles (**Fig. 1I, J**), suggest that 8-week CUMS has not yet caused overt ovarian insufficiency or follicle depletion. Instead, these changes likely reflect a transient compensatory state.

**Figure 2.**
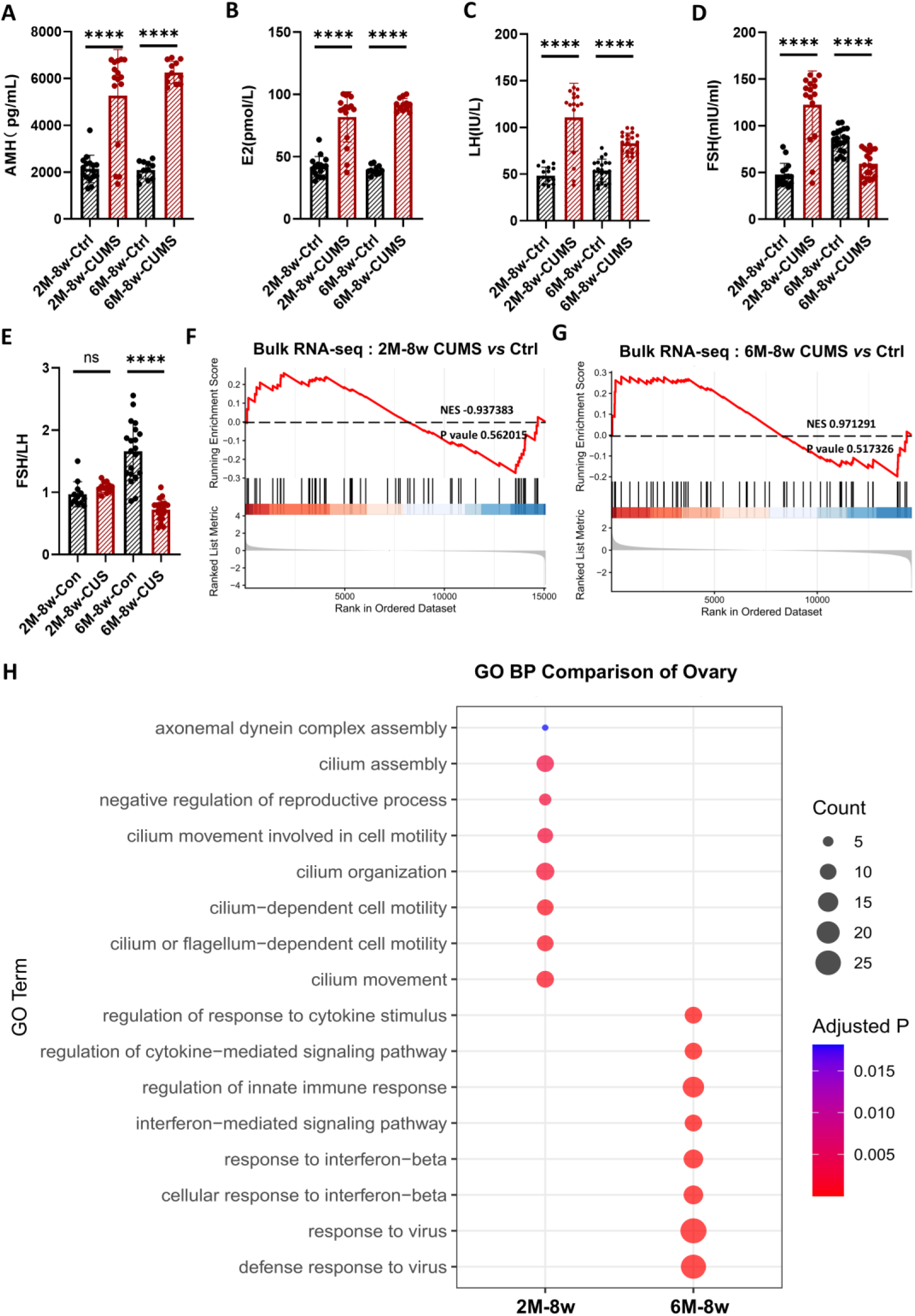
Effects of 8-week CUMS on HPO axis hormones and transcriptomic responses in reproductively young and middle-aged mice. **(A)** Serum AMH levels. **(B)** Serum E2 levels. **(C)** Serum LH levels. **(D)** Serum FSH levels. **(E)** Serum FSH/LH ratio in mice. **(F, G)** GSEA of the GO term “response to estrogen” in the pituitary of the reproductively young group (F) and middle-aged group (G). **(H)** GO analysis of ovarian biological processes. Statistical analysis was performed using unpaired Student’s t-test for two-group comparisons, and one-way ANOVA followed by Tukey’s multiple comparisons test for multi-group comparisons. Data are presented as mean ± SEM. Statistical significance is indicated as ns not significant, * p < 0.05, ** p < 0.01, *** p < 0.001.

In contrast to these consistent elevations, FSH levels displayed a striking age-dependent divergency. While FSH increased significantly in the reproductively young group, it decreased significantly in the reproductively middle-aged group **(Fig. 2D)**. This divergence points to distinct regulatory mechanisms. The decline of FSH in middle-aged mice can be primarily attributed to negative feedback from the elevated E2. In contrast, the sustained elevation of FSH in the reproductively young mice suggests that, at this stage, the pituitary remains predominantly responsive to hypothalamic GnRH signals, while ovarian feedback mechanisms are not yet fully established. Notably, CUMS induced a significant increase of the FSH/LH ratio in reproductively young mice, but a significant decrease in reproductively middle-aged mice **(Fig. 2E)**. These ratio changes strongly support our conclusion that young mice are primarily in a state of pituitary-driven excessive follicle activation, whereas middle-aged mice have engaged an ovarian negative feedback-mediated protective adaptive mechanism.

### 4. Age-dependent ovarian and pituitary transcriptomic responses to 8-week CUMS

To investigate the impact of 8-week CUMS on ovarian and pituitary transcriptional profiles, RNA-seq was performed on samples from both age groups of mice. In the ovaries, principal component analysis (PCA) revealed clear separation between CUMS-exposed and control samples in both reproductively young **(Fig. S2A)** and middle-aged mice **(Fig. S2D)**. Differential expression analysis identified 57 upregulated and 101 downregulated genes in young ovaries exposed to CUMS **(Fig. S2B)**, and 98 upregulated and 61 downregulated genes in middle-aged ovaries exposed to CUMS **(Fig. S2E)**. In the pituitary, PCA also demonstrated clear segregation between CUMS and control samples in the reproductively middle-aged group **(Fig. S3 D)**, whereas no obvious separation was observed in the reproductively young group **(Fig. S3A)**, with 16 upregulated and 8 downregulated genes in young mice exposed to CUMS **(Fig. S3B)**, and 67 upregulated and 63 downregulated genes in middle-aged mice exposed to CUMS **(Fig. S3E)**.

Building on the transcriptomic differences described above, we next sought to elucidate the mechanisms underlying age-dependent differences in FSH **(Fig. 2D)** secretion by assessing pituitary sensitivity to E2 feedback via GSEA of the “response to estrogen” pathway. Following 8 weeks of CUMS, pituitaries from reproductively young mice exhibited downregulation of E2-responsive negative feedback pathways **(NES =** -**0.937383; Fig. 2E)**, whereas those from reproductively middle-aged mice retained responsiveness to estrogen-mediated regulation **(NES = 0.971291; Fig. 2F)**.

GO biological process enrichment analysis revealed that differentially expressed genes (DEGs) in the reproductively young group were predominantly enriched in cilium-related pathways (e.g., “cilium assembly”, “axonemal dynein complex assembly”). In contrast, DEGs in the reproductively middle-aged group were mainly enriched in immune-related pathways (e.g., “regulation of cytokine-mediated signaling response”, “regulation of innate immune responses”) **(Fig. 2H, S2C, S2F)**. These results indicate that 2-month-old mice respond to CUMS with alterations in tissue structural plasticity and ciliary function. In contrast, 6-month-old mice exhibit pronounced immune activation and inflammatory signaling, mirroring early-phase immune activation and inflammatory manifestations reported in psychologically stressed women(18). The distinct biological pathway enrichments observed between the two groups reveal age-dependent pathophysiological divergence in in ovarian stress responses.

### 5. 12-week CUMS induced HPA axis exhaustion and premature ovarian insufficiency in an age-independent manner

Since 8-week CUMS did not induce ovarian atrophy in reproductively middle-aged mice, we extended the CUMS protocol to 12 weeks **(Fig. 3A)**. A significant reduction in body weight remained evident in both reproductively young and middle-aged mice **(Fig. S1D, E)** In contrast to the significant upregulation of circulating corticosterone levels following 8-week CUMS **(Fig. 1B)**, 12-week CUMS led to a significant downregulation of circulating corticosterone levels in both age groups **(Fig. 3B)**, indicating a transition of the HPA axis from compensatory activation during early stress exposure to functional exhaustion or suppression under sustained stress. We also monitored the estrous cycle daily for 14 consecutive days during the modeling period. In both age groups, CUMS significantly prolonged the estrous cycle and caused irregularities in over 80% of mice **(Fig. 3D-G)**.

**Figure 3.**
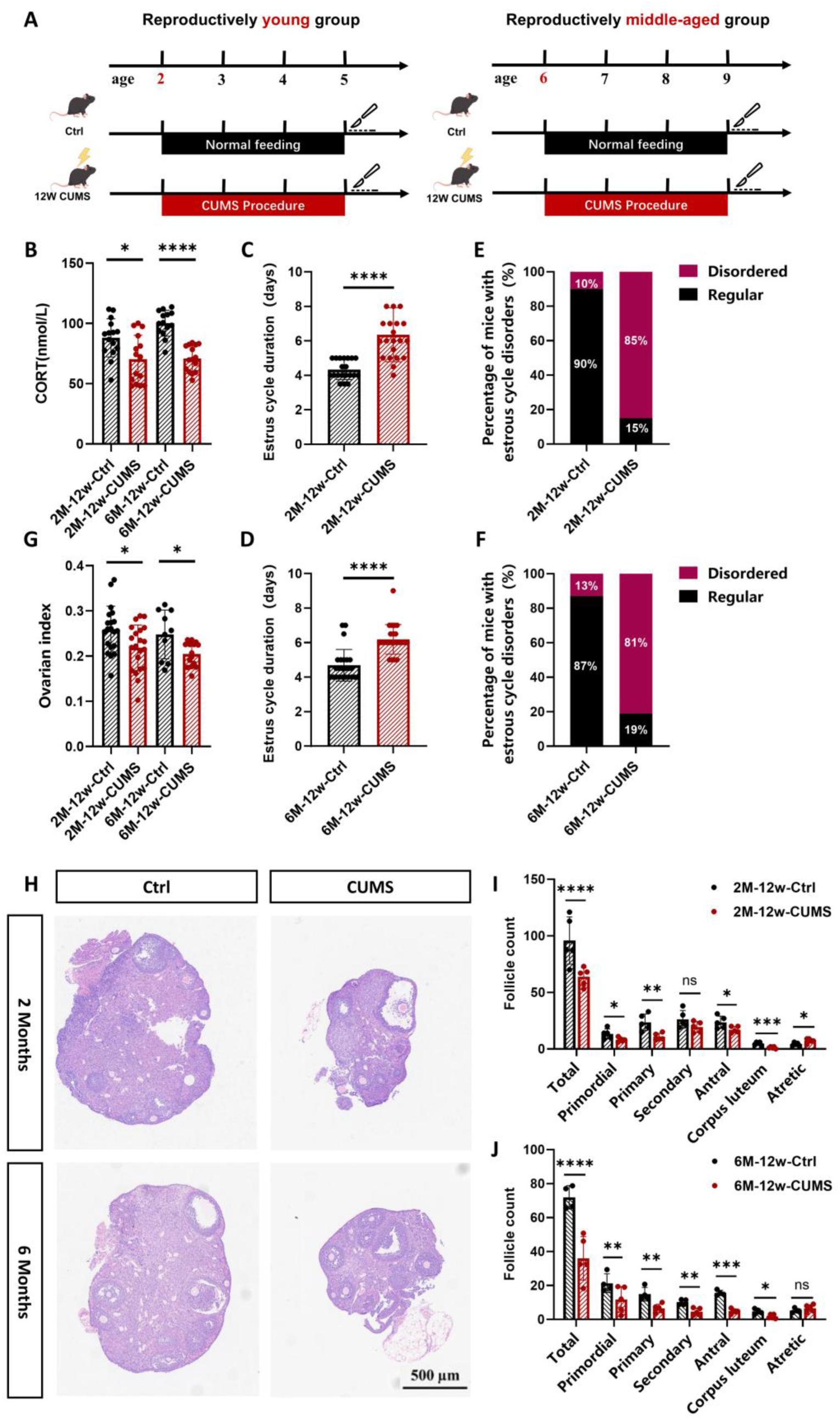
Effects of 12-week CUMS on estrous cyclicity and ovarian reserve in reproductively young and middle-aged mice. **(A)** Schematic diagram of the experimental design. **(B)** Serum corticosterone CORT levels in mice. **(C, D)** Estrus duration in the reproductively young group (C) and middle-aged group (D). **(E,F)** Proportion of mice with estrous cycle irregularities in the reproductively young group (E) and middle-aged group (F). **(G)** Ovarian index of mice. **(H)** HE staining of ovarian follicles. **(I,J)** Counts of different stages of follicles in the reproductively young group (I) and middle-aged group (J) Statistical analysis was performed using unpaired Student’s t-test for two-group comparisons, and one-way ANOVA followed by Tukey’s multiple comparisons test for multi-group comparisons. Data are presented as mean ± SEM. Statistical significance is indicated as ns not significant, * p < 0.05, ** p < 0.01, *** p < 0.001, **** p < 0.0001.

Anatomical assessment of the ovaries revealed a marked decrease in both ovarian weight **(Fig. S1F)** and ovarian index **(Fig. 3G)** in the two age groups. Histological evaluation using H&E staining further demonstrated substantial reductions in follicle numbers across all developmental stages in both reproductively young and middle-aged mice after CUMS exposure **(Fig. 3H-J)**. Together, these findings indicate that 12 weeks of chronic stress accelerates follicle depletion and drives the ovary toward premature insufficiency in both reproductively young and middle-aged mice.

### 6. Age-dependent changes of HPO axis in responses to 12-week CUMS

To further investigate the differential responses of reproductively young and middle-aged group mice to 12-week CUMS, we measured hormone levels associated with the HPO axis. The results showed that both groups displayed significantly decreased AMH and E2 levels, which is consistent with the hormonal profile observed during ovarian insufficiency (**Fig. 4A, B**). LH level was significantly decreased in reproductively middle-aged group (**Fig. 4C)**, which is consistent with the decreased CORT levels suggesting diminished hypothalamic responsiveness to prolonged CUMS. However, reproductively young mice failed to establish normal LH regulation under stress and showed significant increased LH level (**Fig. 4C)**. In addition, both groups exhibited a pronounced reduction in FSH level and FSH/LH (**Fig. 4D, E)**.

**Figure 4.**
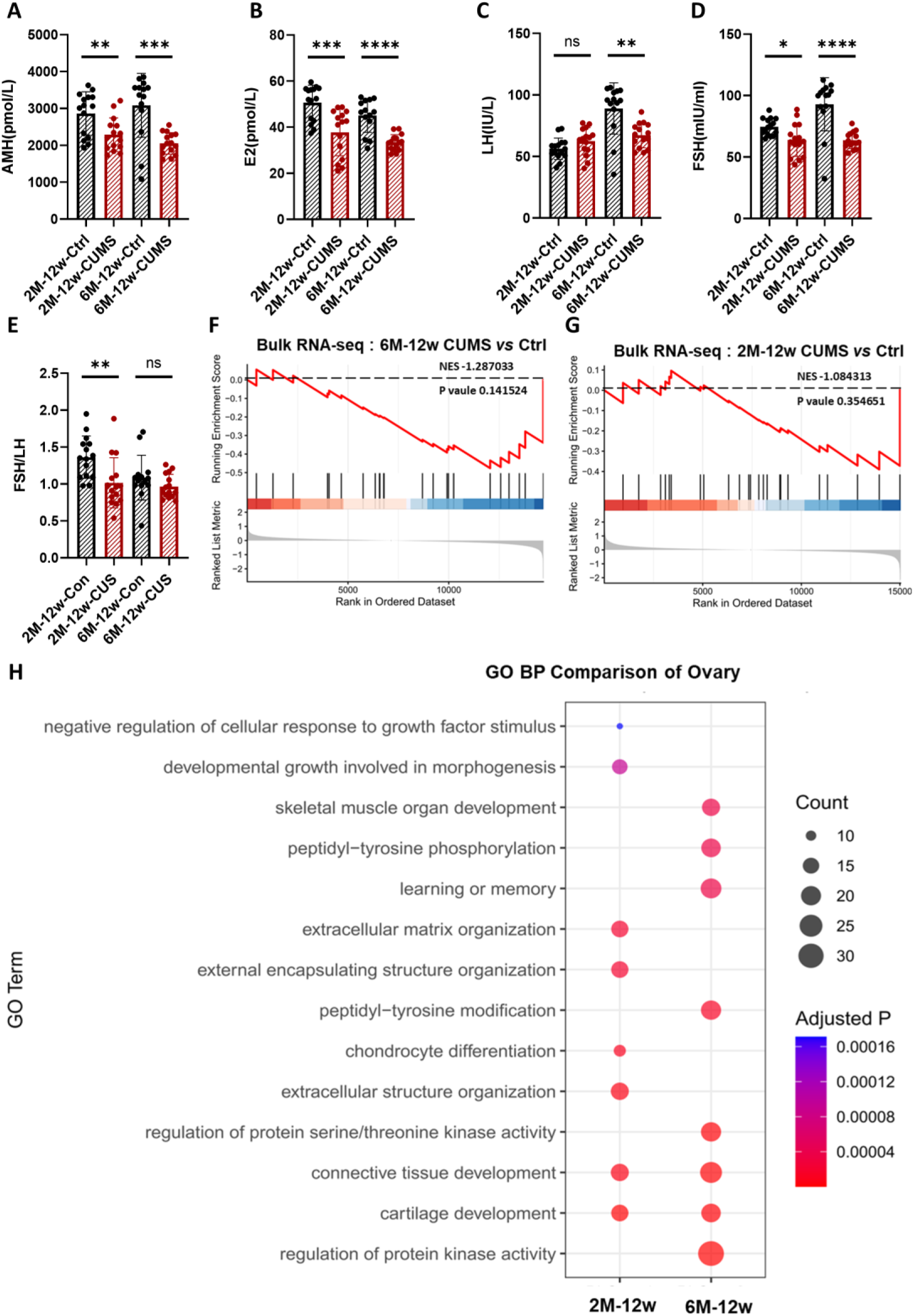
Effects of 12-week CUMS on HPO axis hormones and transcriptomic responses in reproductively young and middle-aged mice. **(A)** Serum AMH levels. **(B)** Serum E2 levels. **(C)** Serum LH levels. **(D)** Serum FSH levels. **(E)** Serum FSH/LH ratio in mice. **(F, G)** GSEA of the GO term “gonadotropin releasing hormone signaling pathway” in the pituitary of the reproductively young group (F) and middle-aged group (G). **(H)** GO analysis of ovarian biological processes. Statistical analysis was performed using unpaired Student’s t-test for two-group comparisons, and one-way ANOVA followed by Tukey’s multiple comparisons test for multi-group comparisons. Data are presented as mean ± SEM. Statistical significance is indicated as ns not significant, * p < 0.05, ** p < 0.01, *** p < 0.001.

To investigate the impact of 12-week CUMS on ovarian and pituitary transcriptional profiles, RNA-seq was performed on samples from both age groups of mice. In the ovaries of reproductively young mice, PCA revealed a clear separation between CUMS-exposed and control samples **(Fig. S4A)**. Differential expression analysis identified 61 upregulated and 223 downregulated genes in response to CUMS **(Fig. S4B)**. In contrast, in the ovaries of reproductively middle-aged mice, PCA demonstrated a clear separation between CUMS and control samples **(Fig. S4D)**, suggesting more pronounced transcriptomic changes. Consistently, differential expression analysis revealed 87 upregulated and 376 downregulated genes following CUMS exposure **(Fig. S4E)**. In the pituitary of reproductively young mice, PCA revealed a clear separation between CUMS-exposed and control samples **(Fig. S5A)**, indicating substantial stress-associated transcriptional remodeling. Differential expression analysis identified 22 upregulated and 32 downregulated genes **(Fig. S5B)**. Conversely, in the pituitary of reproductively middle-aged mice, PCA showed extensive overlap between CUMS and control samples, consistent with relatively moderate transcriptomic changes **(Fig. S5D)**. Correspondingly, only 9 upregulated and 14 downregulated genes were identified in response to CUMS **(Fig. S5E)**.

Interestingly, GSEA revealed significant suppression of gonadotropin-secreting pathways in the pituitary of reproductively young **(Fig. 4F)** and middle-aged **(Fig. 4G)** mice, consistent with the observed decline in LH or FSH levels, since reduced hypothalamic stimulatory input would be expected to result in decreased gonadotrophin secretion. We next performed GO biological process enrichment analysis to identify the key biological processes associated with the differentially expressed genes. In the reproductively young group, differential genes were significantly enriched in pathways related to “extracellular matrix organization”, “external encapsulating structure organization” and “chondrocyte differentiation”, suggesting that the tissue in young mice is predominantly undergoing structural remodeling and matrix maintenance, reflecting an active and orderly extracellular environment. In contrast, the reproductively middle-aged group showed “significant activation of protein kinase–related signaling”, indicating more active signal transduction under chronic stress **(Fig. 4H, S4C, S4F)**. These distinct pathway enrichments further support an age-dependent pathophysiological divergence under chronic stress.

## Discussion

This study systematically compared the characteristics of HPO axis endocrine regulation and ovarian transcriptomic changes in reproductively young group (2-month-old) and middle-aged group (6-month-old) mice following 8-week and 12-week CUMS exposure, At the 8-week time point, we observed that 2-month-old and 6-month-old mice exhibited a reduction in primordial follicles and an increased number of primary follicles, accompanied by elevated levels of AMH, E2, and LH, as well as a corresponding decrease in FSH levels. Unlike 6-month-old mice, 2-month-old mice exhibited impaired pituitary responsiveness to estradiol negative feedback, manifested as dysregulated FSH secretion, suggesting a reduced tolerance to CUMS exposure in younger mice. Following 12-week CUMS exposure, in both 6-month-old and 2-month-old mice, ovarian size and follicle numbers at all stages were significantly reduced, along with decreased AMH, E2, and FSH levels, while LH levels differed between the two age groups. Due to prolonged CUMS exposure, 2-month-old mice failed to establish normal LH regulation, and their LH levels remained persistently elevated. Transcriptomic analyses revealed age-dependent stress responses: ovaries of 2-month-old mice displayed marked activation of inflammatory and immune-related pathways, whereas 6-month-old mice showed sustained upregulation of protein kinase-related signaling networks.

Our results indicate that reproductively young and middle-aged mice adopt entirely distinct endocrine regulatory strategies and aging trajectories in response to stress. Notably, the CUMS model established using reproductively middle-aged mice exhibits higher clinical relevance in simulating the pathological mechanisms of POI in adult women. These findings provide a more physiologically relevant animal model for studying stress-associated reproductive aging, highlight the clear age-dependence of stress-induced endocrine responses, and for the first time describe age-specific differences in stress-induced ovarian aging, offering important insights into reproductive differences across different female age groups.

The advantages of the middle-aged unpredictable stress model are supported by extensive clinical and animal studies, which indicate that the early phase of chronic stress is primarily characterized by immune system activation, including elevated pro-inflammatory factors such as IL-6, TNF-α, and IL-1β, accompanied by enhanced peripheral and central immune responses (19). This inflammation-driven disruption of homeostasis has been considered an early biological hallmark of depression, anxiety, and various stress-related disorders (20). After prolonged stress exposure, the organism gradually enters a stage of impaired neural plasticity and dysregulated cell signaling, characterized by sustained activation of key pathways such as MAPK/ERK, potentially driving profound remodeling of neural structures, synapses, and energy metabolism, ultimately leading to reprogramming of neuro–endocrine–signaling networks (21–23). In the present study, we observed that 6-month-old mice predominantly exhibited upregulation of immune-related pathways after 8 weeks of CUMS, whereas with prolonged stress up to 12 weeks, the major changes shifted toward enhanced protein kinase activity and cell signaling pathways. This time-dependent transition from “immune activation” to “signal network remodeling” closely mirrors the biological progression of long-term psychological stress in humans.

Under 8-week CUMS conditions, both reproductively young group and middle-aged group mice exhibited elevated E2 and AMH levels, but the regulatory patterns of FSH and LH differed significantly. In young mice, a “feedback mismatch” phenomenon was observed, with E2 increasing while FSH remained elevated, suggesting that the negative feedback loop of the HPO axis had not yet fully matured. During puberty and early adulthood, the hypothalamus–pituitary axis remains in a developing state with respect to sex hormone feedback, and this age-specific feedback instability has been previously documented (24). Consequently, under stress, the young pituitary preferentially responds to upstream GnRH signals rather than ovarian-derived negative feedback, resulting in abnormally elevated FSH levels.

This study found that ovarian aging trajectories under chronic stress are markedly age-dependent. In 2-month-old mice, stress is primarily buffered through mechanisms related to “ciliary structure”, “axoneme assembly”, and “tissue plasticity”, representing a “plasticity-type aging trajectory” that maintains early follicle stability via structural remodeling. In contrast, 6-month-old mice tend to activate cytokine, innate immune, and inflammatory pathways, exhibiting a typical “immune-driven aging trajectory.” With prolonged stress, these differences are further amplified: reproductively young ovaries transition from ciliary functional remodeling to extracellular matrix restructuring, whereas reproductively middle-aged ovaries enter deeper stages of aberrant protein kinase and phosphorylation signaling. Overall, reproductively young ovaries maintain function through structural adaptation, while reproductively middle-aged ovaries progress rapidly toward inflammation-associated aging via immune activation. Future studies could integrate single-cell omics, spatial transcriptomics, and multi-omics approaches, while leveraging public single-cell resources (25–27) to improve cell-type annotation and cross-study comparison and to further dissect cell type-specific stress responses and long-term regulatory networks across different ages. Additionally, potential interventions targeting inflammatory responses or senescent cells could be explored to assess their impact on ovarian function and stress sensitivity in reproductively middle-aged ovaries, providing experimental evidence for female reproductive health interventions.

Accumulating evidence has identified chronic psychological stress as an important risk factor for impaired female reproductive health, particularly during midlife—a critical transitional period characterized by the gradual decline of reproductive function. Psychological stress has been shown to disrupt multiple systemic regulatory pathways, including endocrine signaling and metabolic homeostasis, thereby adversely affecting key reproductive processes such as oocyte development and maturation, fertilization, embryo implantation, prregnancy maintenance, and parturition outcomes(8,28–31). Despite growing recognition of the association between stress and reproductive dysfunction, the mechanisms by which chronic stress accelerates reproductive aging during midlife remain poorly understood. Elucidating the stress-responsive regulatory networks governing ovarian function in this vulnerable period is therefore of substantial scientific and clinical significance. A deeper understanding of the plasticity of stress-associated reproductive aging and its potential reversibility may provide critical theoretical support for the development of targeted interventions aimed at delaying or mitigating stress-induced reproductive decline, addressing an increasingly pressing public health challenge.

## Conclusion

This study systematically compared the endocrine regulation of the HPO axis and transcriptomic alterations in 2-month-old (reproductively young group) and 6-month-old (reproductively middle-aged group) mice following 8-week and 12-week CUMS exposure, uncovering pronounced age-dependent divergences in ovarian response to stress. Our findings demonstrate that the CUMS model in reproductively middle-aged mice more faithfully recapitulates the biological processes underlying chronic psychological stress–induced POI in women, underscoring its superior clinical relevance. Notably, reproductively young and middle-aged mice engage fundamentally distinct endocrine regulatory strategies and aging trajectories when subjected to stress. These insights further suggest that age is a critical determinant of stress susceptibility and endocrine adaptability, providing an important conceptual basis for future age-specific investigations into reproductive and endocrine aging. Although transcriptomic analyses provided insights into potential mechanisms underlying differential hormone secretion, these findings remain correlative in nature, indicating associations rather than causality, and require further functional validation to establish causal relationships.

## List of Abbreviations

AMH: Anti-Müllerian Hormone
CORT: Corticosterone
CUMS: Chronic Unpredictable Mild Stress
DEGs: Differentially Expressed Genes
E₂: Estradiol
FDR: False Discovery Rate
FSH: Follicle-Stimulating Hormone
GnRH: Gonadotropin-Releasing Hormone
GO: Gene Ontology
GSEA: Gene Set Enrichment Analysis
HPA: Hypothalamic–Pituitary–Adrenal Axis
HPO: Hypothalamic–Pituitary–Ovarian Axis
LH: Luteinizing Hormone
NES: Normalized Enrichment Score
PCA: Principal Component Analysis
PFA: Paraformaldehyde
POI: Premature Ovarian Insufficiency
ROS: Reactive Oxygen Species
SD: Standard Deviation

## Acknowledgements

We thank the volunteers for participating in this study.

## Author contribution statement

Conceptualization: Jian Li, Zelan Hu,Yudong Wang

methodology: Tiannuo Yang, Shuqin Zhang

software: Tiannuo Yang

validation: Danqi Liu,Shuqin Zhang, Yiqing Ma, Shangxuan Liu

formal analysis: Laijia LI, Hanwen Zhang, Kunpeng Zhou

resources: Jian Li, Zelan Hu, Furui Jin

data curation: Yufeng Han

writing—original draft preparation: Tiannuo Yang, Danqi Liu

All authors have read and agreed to the published version of the manuscript.

## Reference

1. Wang, X., Wang, L. and Xiang, W. (2023) Mechanisms of ovarian aging in women: a review. Journal of Ovarian Research, 16, 67.

2. Zhu, Z., Xu, W. and Liu, L. (2022) Ovarian aging: mechanisms and intervention strategies. Medicinal Research Reviews, 2, 590–610.

3. Wu, C., Chen, D., Stout, M.B., Wu, M. and Wang, S. (2025) Hallmarks of ovarian aging. Trends in Endocrinology and Metabolism, 36, 418–439.

4. European Society for Human, R., Embryology Guideline Group on, P.O.I., Webber, L., Davies, M., Anderson, R., Bartlett, J., Braat, D., Cartwright, B., Cifkova, R., de Muinck Keizer-Schrama, S., et al. (2016) ESHRE Guideline: management of women with premature ovarian insufficiency. Human Reproduction, 31, 926–937.

5. Kala, M. and Nivsarkar, M. (2016) Role of cortisol and superoxide dismutase in psychological stress induced anovulation. General and Comparative Endocrinology, 225, 117–124.

6. 6. Smits, M.A.J., Schomakers, B.V., van Weeghel, M., Wever, E.J.M., Wust, R.C.I., Dijk, F., Janssens, G.E., Goddijn, M., Mastenbroek, S., Houtkooper, R.H., et al. (2023) Human ovarian aging is characterized by oxidative damage and mitochondrial dysfunction. Human Reproduction, 38, 2208–2220.

7. Yan, F., Zhao, Q., Li, Y., Zheng, Z., Kong, X., Shu, C., Liu, Y. and Shi, Y. (2022) The role of oxidative stress in ovarian aging: a review. Journal of Ovarian Research, 15, 100.

8. Hu, Y., Wang, W., Ma, W., Wang, W., Ren, W., Wang, S., Fu, F. and Li, Y. (2025) Impact of psychological stress on ovarian function: Insights, mechanisms and intervention strategies (Review). International Journal of Molecular Medicine, 55.

9. Gao, L., Zhao, F., Zhang, Y., Wang, W. and Cao, Q. (2020) Diminished ovarian reserve induced by chronic unpredictable stress in C57BL/6 mice. Gynecological Endocrinology, 36, 49–54.

10. Xiang, Y., Jiang, L., Gou, J., Sun, Y., Zhang, D., Xin, X., Song, Z. and Huang, J. (2022) Chronic unpredictable mild stress-induced mouse ovarian insufficiency by interrupting lipid homeostasis in the ovary. Frontiers in Cell and Developmental Biology, 10, 933674.

11. Ma, J., Wang, L., Yang, D., Luo, J., Gao, J., Wang, J., Guo, H., Li, J., Wang, F., Wu, J. et al. (2024) Chronic stress causes ovarian fibrosis to impair female fertility in mice. Cellular Signalling, 122, 111334.

12. Ding, S.M., Shi, L.G., Xing, F., Cui, S.S., Cheng, H.R., Liu, Y., Ji, D.M., Liang, D., Cao, Y.X. and Liu, Y.J. (2024) Melatonin Protects Against Mitochondrial Dyshomeostasis and Ovarian Damage Caused by Chronic Unpredictable Mild Stress Through the eIF2alpha-AFT4 Signaling Pathway in Mice. Reproductive Sciences, 31, 3191–3201.

13. Willner, P. (2017) The chronic mild stress (CMS) model of depression: History, evaluation and usage. Neurobiology of Stress, 6, 78–93.

14. Zhu, Y., Sun, H., Gao, T., Hou, S., Li, Y., Xu, Y., Zhang, Q. and Feng, D. (2025) Ovarian remodeling and aging-related chronic inflammation and fibrosis in the mammalian ovary. J Ovarian Res, 18, 133.

15. Dipali, S.S., King, C.D., Rose, J.P., Burdette, J.E., Campisi, J., Schilling, B. and Duncan, F.E. (2023) Proteomic quantification of native and ECM-enriched mouse ovaries reveals an age-dependent fibro-inflammatory signature. Aging (Albany NY), 15, 10821–10855.

16. Balough, J.L., Dipali, S.S., Velez, K., Kumar, T.R. and Duncan, F.E. (2024) Hallmarks of female reproductive aging in physiologic aging mice. Nature Aging, 4, 1711–1730.

17. Sievert, L.L., Jaff, N. and Woods, N.F. (2018) Stress and midlife women’s health. Women’s Midlife Health, 4, 4.

18. Endrighi, R., Hamer, M. and Steptoe, A. (2016) Post-menopausal Women Exhibit Greater Interleukin-6 Responses to Mental Stress Than Older Men. Ann Behav Med, 50, 564–571.

19. Hodes, G.E., Kana, V., Menard, C., Merad, M. and Russo, S.J. (2015) Neuroimmune mechanisms of depression. Nature Neuroscience, 18, 1386–1393.

20. Liu, Y.Z., Wang, Y.X. and Jiang, C.L. (2017) Inflammation: The Common Pathway of Stress-Related Diseases. Frontiers in Human Neuroscience, 11, 316.

21. Wohleb, E.S., Powell, N.D., Godbout, J.P. and Sheridan, J.F. (2013) Stress-induced recruitment of bone marrow-derived monocytes to the brain promotes anxiety-like behavior. The Journal of Neuroscience, 33, 13820–13833.

22. Miller, G.E., Rohleder, N. and Cole, S.W. (2009) Chronic interpersonal stress predicts activation of pro- and anti-inflammatory signaling pathways 6 months later. Psychosomatic Medicine, 71, 57–62.

23. Pace, T.W., Mletzko, T.C., Alagbe, O., Musselman, D.L., Nemeroff, C.B., Miller, A.H. and Heim, C.M. (2006) Increased stress-induced inflammatory responses in male patients with major depression and increased early life stress. American Journal of Psychiatry, 163, 1630–1633.

24. Kurian, J.R. and Terasawa, E. (2013) Epigenetic control of gonadotropin releasing hormone neurons. Frontiers in Endocrinology, 4, 61.

25. Zhang, J., Jia, S., Zheng, Z., Cao, L., Zhou, J. and Fu, X. (2025) A multi-omic single-cell landscape of the aging mouse ovary. GeroScience, 47, 4485–4498.

26. Madrigal, P., Thanki, A.S., Fexova, S., Yu, I.D., Chatzigeorgiou, A., Zucchi, I., Marugan Calles, J.C., Vilmovsky, L., Khen, A., Zhao, L. et al. (2026) Expression Atlas in 2026: enabling FAIR and open expression data through community collaboration and integration. Nucleic Acids Research, 54, D147–D157.

27. Lian, X., Zhang, Y., Zhou, Y., Sun, X., Huang, S., Dai, H., Han, L. and Zhu, F. (2024) SingPro: a knowledge base providing single-cell proteomic data. Nucleic Acids Research, 52, D552–D561.

28. Sun, J., Fan, Y., Guo, Y., Pan, H., Zhang, C., Mao, G., Huang, Y., Li, B., Gu, T., Wang, L. et al. (2022) Chronic and Cumulative Adverse Life Events in Women with Primary Ovarian Insufficiency: An Exploratory Qualitative Study. Frontiers in Endocrinology, 13, 856044.

29. Zhai, Q.Y., Wang, J.J., Tian, Y., Liu, X. and Song, Z. (2020) Review of psychological stress on oocyte and early embryonic development in female mice. Reproductive Biology and Endocrinology, 18, 101.

30. Prasad, S., Tiwari, M., Pandey, A.N., Shrivastav, T.G. and Chaube, S.K. (2016) Impact of stress on oocyte quality and reproductive outcome. Journal of Biomedical Science, 23, 36.

31. Vrekoussis, T., Kalantaridou, S.N., Mastorakos, G., Zoumakis, E., Makrigiannakis, A., Syrrou, M., Lavasidis, L.G., Relakis, K. and Chrousos, G.P. (2010) The role of stress in female reproduction and pregnancy: an update. Annals of the New York Academy of Sciences, 1205, 69–75.

