## Supplemental File 1 for "Distinct HPO axis responses and ovarian aging trajectories to chronic unpredictable mild stress in reproductively young versus middle-aged female mice"

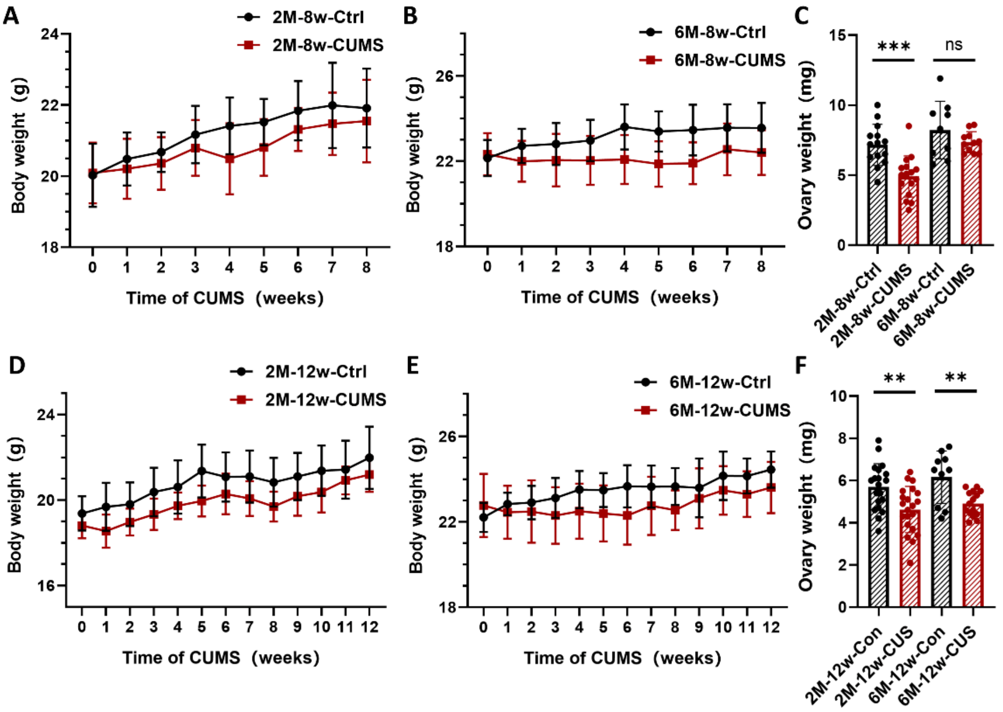
A
2M-8w-Ctrl
2M-8w-CUMS
Body weight (g)
Time of CUMS (weeks)
B
6M-8w-Ctrl
6M-8w-CUMS
Body weight (g)
Time of CUMS (weeks)
C
***
ns
Ovary weight (mg)
2M-8w-Ctrl
2M-8w-CUMS
6M-8w-Ctrl
6M-8w-CUMS
D
2M-12w-Ctrl
2M-12w-CUMS
Body weight (g)
Time of CUMS (weeks)
E
6M-12w-Ctrl
6M-12w-CUMS
Body weight (g)
Time of CUMS (weeks)
F
**
**
Ovary weight (mg)
2M-12w-Con
2M-12w-CUS
6M-12w-Con
6M-12w-CUS

**Supplementary Figure S1**.

**Effects of CUMS on body weight and ovarian weight in mice. (A)** Body weight changes in the reproductively young control group (n = 16) and CUMS group (n = 16) following 8-week CUMS intervention. **(B)** Body weight changes in the reproductively middle-aged control group (n = 11) and CUMS Group (n = 12) following 8-week CUMS intervention. **(C)** Ovary weight changes in mice following 8-week CUMS intervention. **(D)** Body weight curves in reproductively young control group (n = 15) and CUMS group (n = 15) following 12-week CUMS intervention. **(E)** Body weight curves in reproductively aged control group (n = 14) and CUMS group (n = 15) following 12-week CUMS intervention. **(F)** Ovary weight changes in mice following 12-week CUMS intervention.Statistical analysis was performed using unpaired Student’s t-test for two-group comparisons, and one-way ANOVA followed by Tukey’s multiple comparisons test for multi-group comparisons. Data are presented as mean ± SEM. Statistical significance is indicated as ns not significant, * $p < 0.05$, ** $p < 0.01$, *** $p < 0.001$.

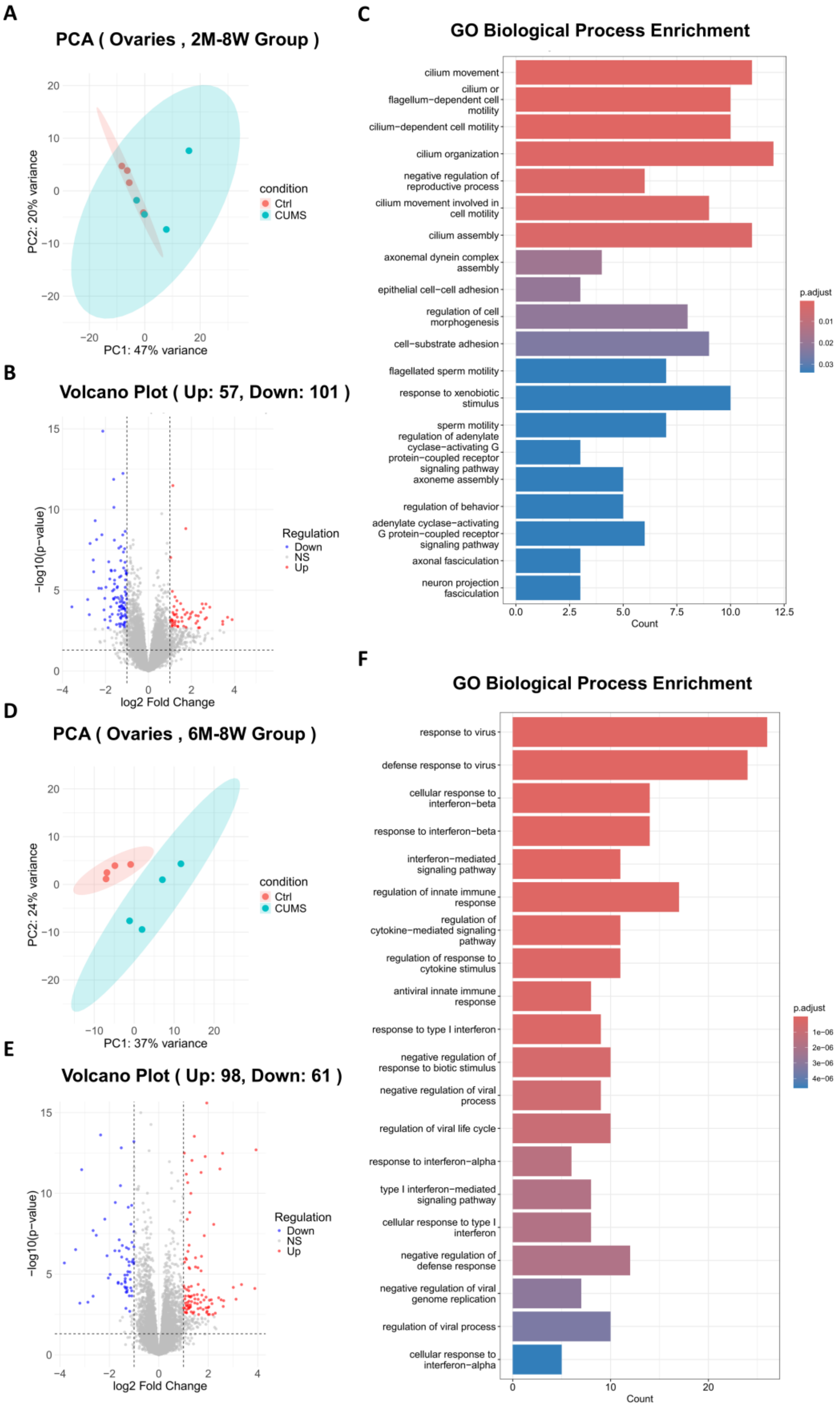
A
PCA ( Ovaries , 2M-8W Group )
PC2: 20% variance
PC1: 47% variance
condition
Ctrl
CUMS
B
Volcano Plot ( Up: 57, Down: 101 )
−log10(p−value)
log2 Fold Change
Regulation
Down
NS
Up
C
GO Biological Process Enrichment
cilium movement
cilium or flagellum−dependent cell motility
cilium−dependent cell motility
cilium organization
negative regulation of reproductive process
cilium movement involved in cell motility
cilium assembly
axonemal dynein complex assembly
epithelial cell−cell adhesion
regulation of cell morphogenesis
cell−substrate adhesion
flagellated sperm motility
response to xenobiotic stimulus
sperm motility
regulation of adenylate cyclase−activating G protein−coupled receptor signaling pathway
axoneme assembly
regulation of behavior
adenylate cyclase−activating G protein−coupled receptor signaling pathway
axonal fasciculation
neuron projection fasciculation
Count
p.adjust
D
PCA ( Ovaries , 6M-8W Group )
PC2: 24% variance
PC1: 37% variance
condition
Ctrl
CUMS
E
Volcano Plot ( Up: 98, Down: 61 )
−log10(p−value)
log2 Fold Change
Regulation
Down
NS
Up
F
GO Biological Process Enrichment
response to virus
defense response to virus
cellular response to interferon−beta
response to interferon−beta
interferon−mediated signaling pathway
regulation of innate immune response
regulation of cytokine−mediated signaling pathway
regulation of response to cytokine stimulus
antiviral innate immune response
response to type I interferon
negative regulation of response to biotic stimulus
negative regulation of viral process
regulation of viral life cycle
response to interferon−alpha
type I interferon−mediated signaling pathway
cellular response to type I interferon
negative regulation of defense response
negative regulation of viral genome replication
regulation of viral process
cellular response to interferon−alpha
Count
p.adjust

**Supplementary Figure S2.**

**Effects of 8-week CUMS on the ovarian transcriptome of mice. (A)** Principal component analysis of the reproductively young group. **(B)** volcano plots of the reproductively young group. **(C)** Gene Ontology enrichmen of the reproductively young group. **(D)** Principal component analysis of the reproductively middle-aged group. **(E)** volcano plots of the reproductively group. **(F)** Gene Ontology enrichmen of the reproductively middle-aged group.

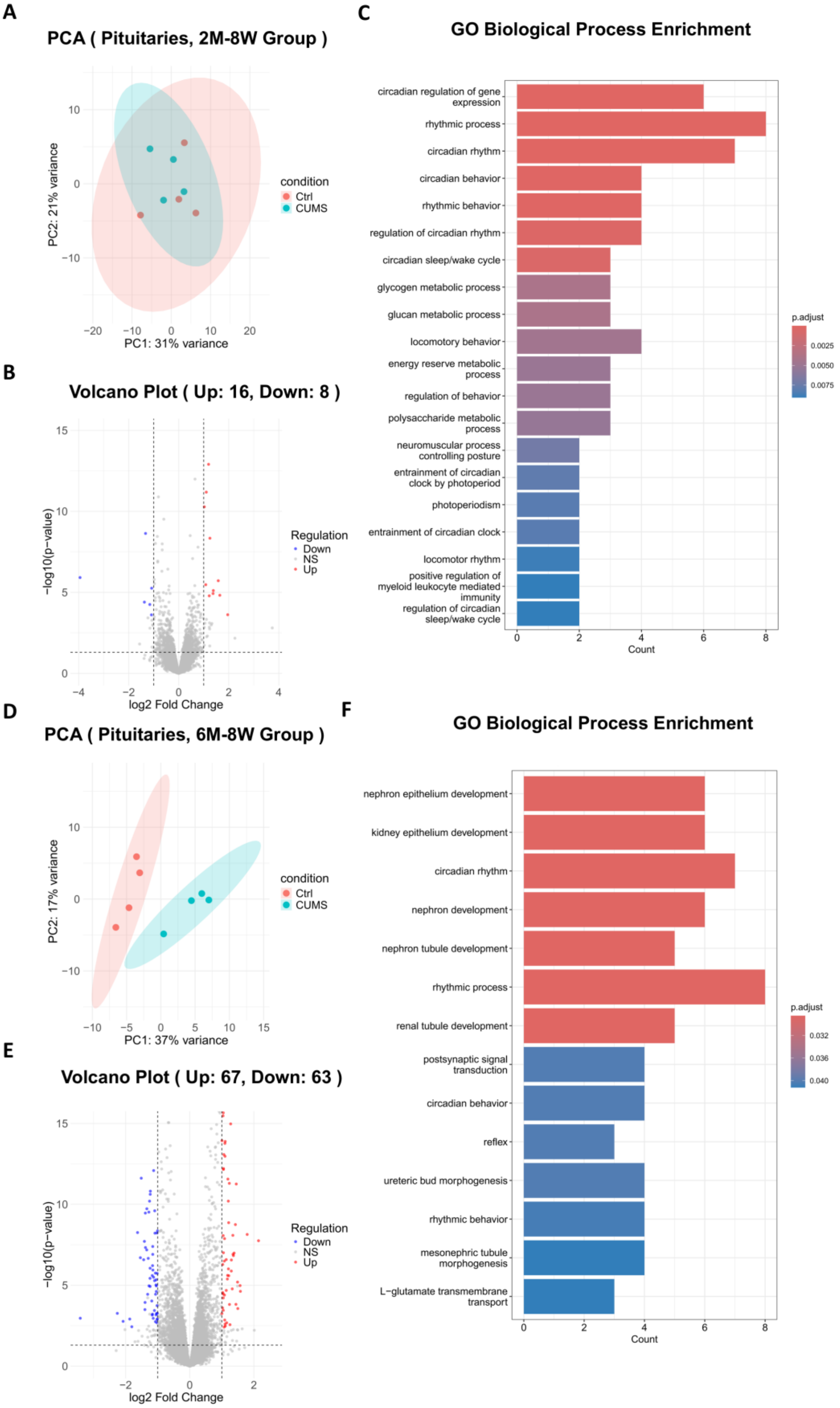
A
PCA ( Pituitaries, 2M-8W Group )
PC2: 21% variance
PC1: 31% variance
condition
Ctrl
CUMS
B
Volcano Plot ( Up: 16, Down: 8 )
−log10(p−value)
log2 Fold Change
Regulation
Down
NS
Up
C
GO Biological Process Enrichment
circadian regulation of gene expression
rhythmic process
circadian rhythm
circadian behavior
rhythmic behavior
regulation of circadian rhythm
circadian sleep/wake cycle
glycogen metabolic process
glucan metabolic process
locomotory behavior
energy reserve metabolic process
regulation of behavior
polysaccharide metabolic process
neuromuscular process controlling posture
entrainment of circadian clock by photoperiod
photoperiodism
entrainment of circadian clock
locomotor rhythm
positive regulation of myeloid leukocyte mediated immunity
regulation of circadian sleep/wake cycle
Count
p.adjust
0.0025
0.0050
0.0075
D
PCA ( Pituitaries, 6M-8W Group )
PC2: 17% variance
PC1: 37% variance
condition
Ctrl
CUMS
E
Volcano Plot ( Up: 67, Down: 63 )
−log10(p−value)
log2 Fold Change
Regulation
Down
NS
Up
F
GO Biological Process Enrichment
nephron epithelium development
kidney epithelium development
circadian rhythm
nephron development
nephron tubule development
rhythmic process
renal tubule development
postsynaptic signal transduction
circadian behavior
reflex
ureteric bud morphogenesis
rhythmic behavior
mesonephric tubule morphogenesis
L−glutamate transmembrane transport
Count
p.adjust
0.032
0.036
0.040

**Supplementary Figure S3.**

**Effects of 8-week CUMS on the pituitary transcriptome of mice. (A)** Principal component analysis of the reproductively young group. **(B)** volcano plots of the reproductively young group. **(C)** Gene Ontology enrichmen of the reproductively young group. **(D)** Principal component analysis of the reproductively middle-aged group. **(E)** volcano plots of the reproductively group. **(F)** Gene Ontology enrichmen of the reproductively middle-aged grop.

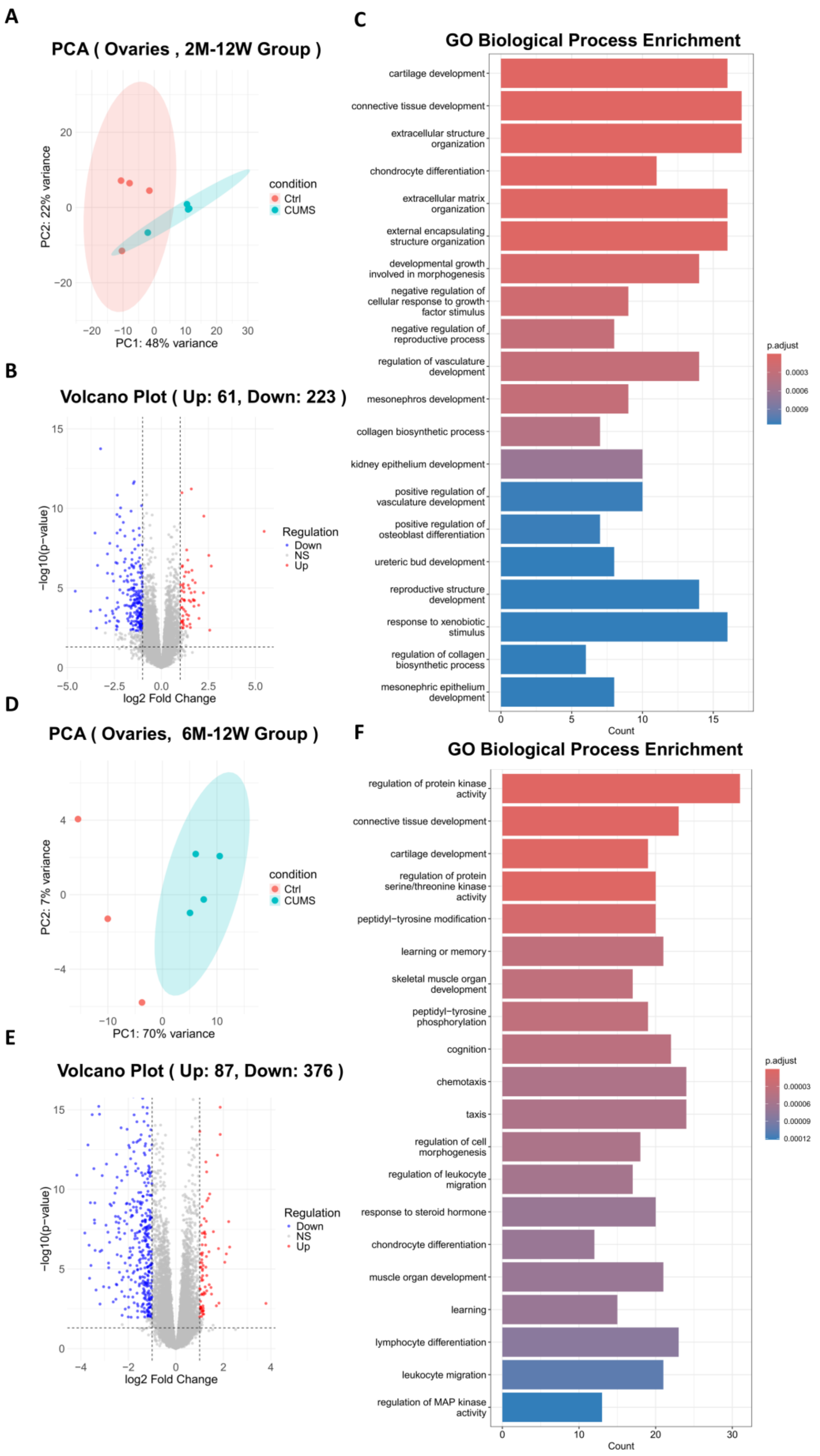
A
PCA ( Ovaries , 2M-12W Group )
PC1: 48% variance
PC2: 22% variance
condition
Ctrl
CUMS
B
Volcano Plot ( Up: 61, Down: 223 )
log2 Fold Change
−log10(p−value)
Regulation
Down
NS
Up
C
GO Biological Process Enrichment
cartilage development
connective tissue development
extracellular structure organization
chondrocyte differentiation
extracellular matrix organization
external encapsulating structure organization
developmental growth involved in morphogenesis
negative regulation of cellular response to growth factor stimulus
negative regulation of reproductive process
regulation of vasculature development
mesonephros development
collagen biosynthetic process
kidney epithelium development
positive regulation of vasculature development
positive regulation of osteoblast differentiation
ureteric bud development
reproductive structure development
response to xenobiotic stimulus
regulation of collagen biosynthetic process
mesonephric epithelium development
Count
p.adjust
D
PCA ( Ovaries, 6M-12W Group )
PC1: 70% variance
PC2: 7% variance
condition
Ctrl
CUMS
E
Volcano Plot ( Up: 87, Down: 376 )
log2 Fold Change
−log10(p−value)
Regulation
Down
NS
Up
F
GO Biological Process Enrichment
regulation of protein kinase activity
connective tissue development
cartilage development
regulation of protein serine/threonine kinase activity
peptidyl−tyrosine modification
learning or memory
skeletal muscle organ development
peptidyl−tyrosine phosphorylation
cognition
chemotaxis
taxis
regulation of cell morphogenesis
regulation of leukocyte migration
response to steroid hormone
chondrocyte differentiation
muscle organ development
learning
lymphocyte differentiation
leukocyte migration
regulation of MAP kinase activity
Count
p.adjust

**Supplementary Figure S4.**

**Effects of 12-week CUMS on the ovarian transcriptome of mice. (A)** Principal component analysis of the reproductively young group. **(B)** volcano plots of the reproductively young group. **(C)** Gene Ontology enrichmen of the reproductively young group. **(D)** Principal component analysis of the reproductively middle-aged group. **(E)** volcano plots of the reproductively group. **(F)** Gene Ontology enrichmen of the reproductively middle-aged group.

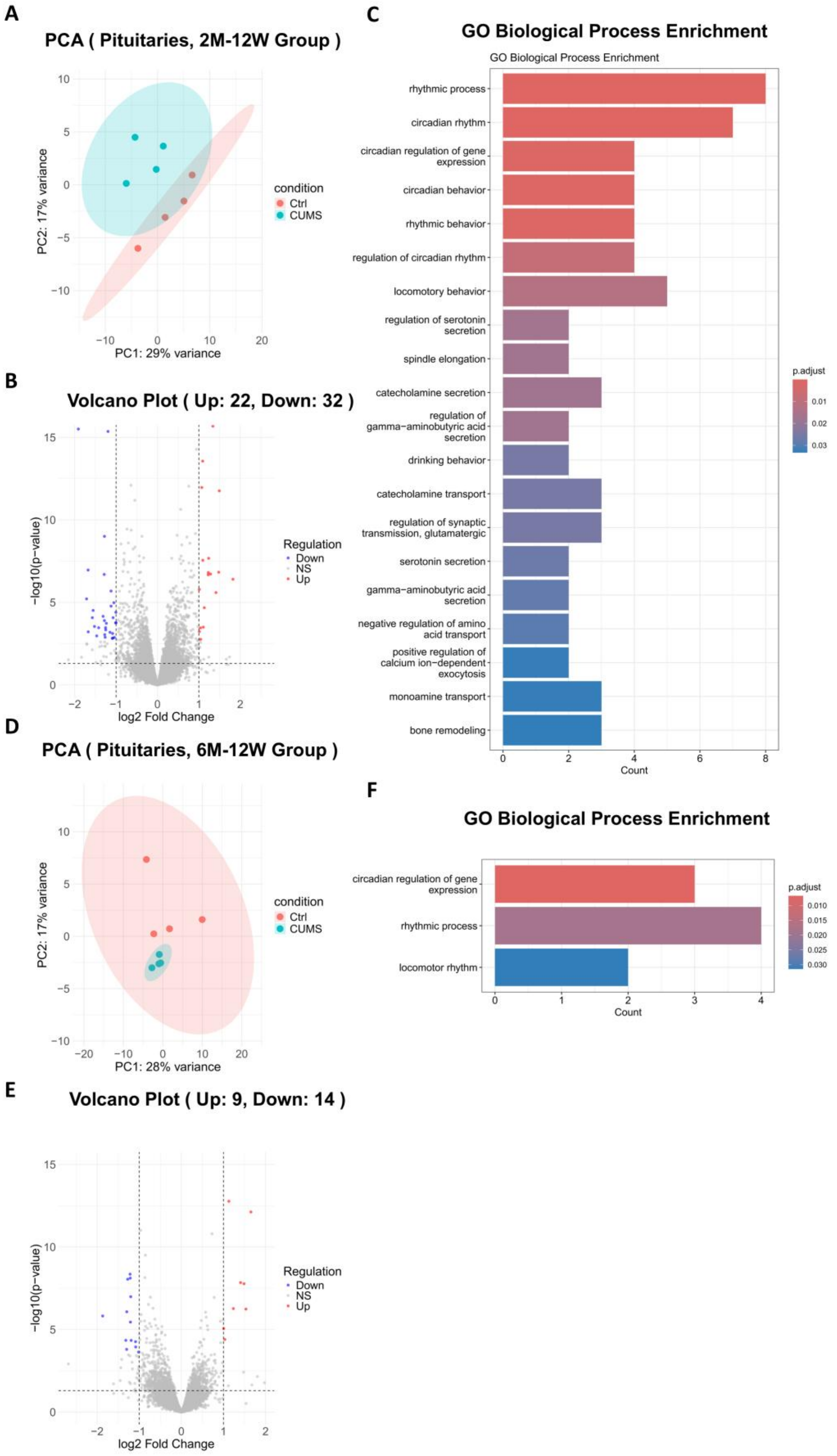
A
PCA ( Pituitaries, 2M-12W Group )
PC2: 17% variance
PC1: 29% variance
condition
Ctrl
CUMS
B
Volcano Plot ( Up: 22, Down: 32 )
−log10(p−value)
log2 Fold Change
Regulation
Down
NS
Up
C
GO Biological Process Enrichment
GO Biological Process Enrichment
rhythmic process
circadian rhythm
circadian regulation of gene expression
circadian behavior
rhythmic behavior
regulation of circadian rhythm
locomotory behavior
regulation of serotonin secretion
spindle elongation
catecholamine secretion
regulation of gamma−aminobutyric acid secretion
drinking behavior
catecholamine transport
regulation of synaptic transmission, glutamatergic
serotonin secretion
gamma−aminobutyric acid secretion
negative regulation of amino acid transport
positive regulation of calcium ion−dependent exocytosis
monoamine transport
bone remodeling
Count
p.adjust
D
PCA ( Pituitaries, 6M-12W Group )
PC2: 17% variance
PC1: 28% variance
condition
Ctrl
CUMS
E
Volcano Plot ( Up: 9, Down: 14 )
−log10(p−value)
log2 Fold Change
Regulation
Down
NS
Up
F
GO Biological Process Enrichment
circadian regulation of gene expression
rhythmic process
locomotor rhythm
Count
p.adjust

**Supplementary Figure S5.**

**Effects of 12-week CUMS on the pituitary transcriptome of mice. (A)** Principal component analysis of the reproductively young group. **(B)** Volcano plots of the reproductively young group. **(C)** Gene Ontology enrichmen of the reproductively young group. **(D)** Principal component analysis of the reproductively middle-aged group. **(E)** volcano plots of the reproductively group. **(F)** Gene Ontology enrichmen of the reproductively middle-aged group.
